# Role of guard-cell ABA in determining maximal stomatal aperture and prompt vapor-pressure-deficit response

**DOI:** 10.1101/218719

**Authors:** Adi Yaaran, Boaz Negin, Menachem Moshelion

## Abstract

Abscisic acid (ABA) is known to be involved in stomatal closure. However, its role in stomatal response to rapid increases in the vapor pressure deficit (VPD) is unclear. To study this issue, we generated guard cell (GC)-specific ABA-insensitive *Arabidopsis* plants (GC-specific *abi1-1*; GCabi). Under normal conditions, the stomatal conductance (g_s_) and apertures of GCabi plants were greater than those of control plants. This supports GC ABA role as limiting maximal stomatal aperture under non-stressful conditions. When there was a rapid increase in VPD (0.15 to 1 kPa), the g_s_ and stomatal apertures of GCabi decreased in a manner similar that observed in the WT control, but different from that observed in WT plants treated with fusicoccin. Low VPD increased the size of the stomatal apertures of the WT, but not of GCabi. We conclude that GC ABA does not play a significant role in the initial, rapid stomatal closure that occurs in response to an increase in VPD, but is important for stomatal adaptation to ambient VPD. We propose a biphasic angiosperm VPD-sensing model that includes an initial passive-hydraulic, ABA-independent phase and a subsequent ABA-dependent steady-state phase in which stomatal behavior is optimized for ambient VPD conditions.

**Highlight:** Guard-cell ABA does not play a significant role in the immediate closure of stomata following an increase in the VPD, but is important for stomatal adaptation to ambient VPD.

## 1. Introduction

Stomata are microscopic pores that allow for controlled gas exchange between a plant and the atmosphere. In early-diverging vascular plants (i.e., ferns), stomatal control displays passive-hydraulic characteristics, such that small decreases in turgor result in rapid reductions in stomatal aperture, which are accompanied by a decrease in the rate of CO_2_ assimilation (Brodribb and McAdam, 2011). The emergence of an abscisic acid (ABA)-dependent stomatal regulation mechanism (approximately 360 million years ago) increased the flexibility of stomatal control (Brodribb and McAdam, 2011). This active-chemical mechanism initiates rapid signal transduction for the depolarization of guard cell (GC) membrane potential, decreased osmotic concentration, turgor loss and reduced stomatal aperture (Daszkowska-Golec and Szarejko, 2013; Munemasa *et al.*, 2015).

Stomatal aperture is known to respond to differences between the vapor concentration within the leaf and the vapor concentration in the air. Mott (1991) showed that GC do not sense relative humidity (RH) directly, but do respond to changes in the transpiration rate. The atmospheric vapor pressure deficit (VPD) serves as the driving force for transpiration, determining the rate at which water is lost from the leaf. An increase in the VPD (a greater difference in vapor concentrations) accelerates the loss of water from the leaf and initiates a reduction in stomatal aperture that prevents excessive water loss and protects the leaf from desiccation. Due to its important role in stomatal regulation, ABA has been considered as a possible key player in the mechanism by which the GC respond to changes in the VPD. McAdam and Brodribb (2016) recently showed that a reduction in leaf turgor can trigger ABA biosynthesis and that increased sensitivity of ABA synthesis to leaf turgor corresponds with a higher stomatal sensitivity to VPD, suggesting that the rapid biosynthesis of ABA in the leaf (~ 10 min) could be responsible for the angiosperms’ stomatal VPD response (McAdam *et al.*, 2015; Sussmilch *et al.*, 2017). Moreover, an increase in GC ABA was measured 15 min after a drop in humidity (i.e., an increase in VPD; Waadt *et al.*, 2014). Indeed, GC were shown to possess the entire ABA biosynthesis pathway (Bauer *et al.*, 2013), which is sufficient for the stomatal response to low RH (Merilo *et al.*, 2017). These last two findings support the hypothesis that GC self-synthesize ABA in response to an increase in VPD. However, other studies have suggested that ABA synthesis is not limited to the GC and that more intense ABA synthesis may take place elsewhere in the leaf (McAdam and Brodribb 2015). Mutants with impaired ABA metabolism (Xie *et al.*, 2006; Okamoto *et al.*, 2008; Merilo *et al.*, 2013; Bauer *et al.*, 2013; McAdam *et al.*, 2015 *Arabidopsis thaliana, Pisum sativum, Solanum lycopersicon*) and signaling (Xie *et al.*, 2006; Ache *et al.*, 2010; Merilo *et al.*, 2013; *Arabidopsis thaliana*) exhibited impaired stomatal responses to rapid changes in VPD. Buckley (2015) recently claimed that the increase in ABA content following an increase in the VPD can fill in a gap in the hydro-active feedback hypothesis, demonstrating how an ultimate mechanism (gene regulation) yields an intermediate signal and a proximate effect (stomatal closure).

There is also other evidence that is not congruent with the hypothesis that ABA is the main cause for the angiosperm stomatal VPD response. In a very recent study, Merilo *et al.* (2017) showed that a broad range of Arabidopsis ABA mutants (mostly ABA-deficient) exhibit a reduction in g_s_ in response to an immediate increase in VPD that is similar or even more intense than that observed for the WT, raising anew the debate regarding the role of ABA in this process. This new evidence corresponds with the work of Assmann *et al.* (2000), which showed that ABA-deficient (*aba1*) and ABA-insensitive (*abi1-1*, *abi2-1*) Arabidopsis mutants have a WT-like stomatal response to VPD, as well as the ABA-independent VPD stomatal closure pathway reported by Yoshida *et al.* (2006) and Merilo *et al.* (2017).

In order to better understand the role of GC ABA in angiosperms’ responses to VPD, we generated, for the first time, GC-specific ABA-insensitive plants (GCabi *Arabidopsis thaliana* plants) using the *abi1-1* mutant gene under the control of a GC-specific promoter, resulting in dominant GC ABA insensitivity against a non-manipulated background. We demonstrate that while GC ABA does play a role in adjusting stomatal aperture to the ambient VPD, it plays no role in sensing rapid changes in ambient VPD, which seems to decrease stomatal conductance and stomatal apertures via a mechanism that is not ABA-dependent.

## 2. Materials and methods

### 2.1. Plant material

Arabidopsis (*Arabidopsis thaliana* ecotype Colombia) lines that express GFP or abi1-1 specifically in guard cells (GCGFP and GCabi lines, respectively) were generated following transformation with GFP or abi1-1 expressed under the KST1 promoter (Müller-Röber *et al.*, 1995) using the floral-dip transformation method (Clough and Bent, 1998). *abi1-1* is a gain-of-function mutation that also has dominant negative features in terms of ABA-sensing (Koornneef *et al.*, 1984; Gosti *et al.*, 1999; Park *et al.*, 2009). Expression of abi1-1 results in ABA insensitivity despite the presence of the WT ABI and other redundant PP2Cs, even when the Arabidopsis gene is expressed in poplar (*Populus* x *canescens* [Ait.] Sm.; Arend *et al.*, 2009) and tomato (*Lycopersicon esculentum* L.; Carrera and Prat, 1998). Independent transgenic lines for each construct were identified.

The Arabidopsis plants were grown in a growth chamber under short-day conditions (10 h light, light intensity of ~150 μmol m−2 s−1) at a controlled temperature of 20– 22°C. Plants were exposed to 50% humidity (VPD = ~1.4 kPa) or covered with clear plastic lid to maintain 90% humidity (VPD = ~0.2 kPa). All plants were grown in potting mix containing (w/w) 30% vermiculite, 30% peat, 20% tuff and 20% perlite (Shacham; Beit Haemek, Israel).

*Cyrtomium falcatum* ferns were obtained from the Givat Brenner Nursery (Israel) and were grown in a tropical greenhouse until they were transferred to the lysimeter system (described below).

### 2.2. Generation of GCabi and GCGFP plants

The *abi1-1* gene from an *abi1-1* plant (Landsberg ecotype) was cloned into the pDONR™ 221 vector (Invitrogen; Waltham, MA USA) and the KST promoter (Müller-Röber *et al.*, 1995) was cloned into pDONRP4P1r using Gateway BP reactions, and later recombined into a pK7M24GW two-fragment destination vector (Karimi *et al.*, 2007) using a Gateway LR reaction, according to the manufacturer’s instructions. The binary *KST:abi1-1* vector was transformed into agrobacterium by electroporation. The *KST:GFP* binary vector was constructed using the same method used to construct the *KST:abi1-1* binary vector, except that the *abi1-1* gene was replaced with the GFP (green fluorescent protein) gene. GCabi mutants were identified through the use of high-resolution melt analysis real-time PCR (Corbett Research Rotor-Gene 6000 cycler; Sydney, Australia) using forward (5TGGTCGGTTTGATCCTCAAT3) and reverse (5TAGCTATCTCCTCCGCCAAA3) primers. DNA of plants suspected to be transgenic was sequenced to confirm the presence of the *abi1-1* snip (G to A at position 539) in the plant.

Comparison of four independent GCabi lines to the WT revealed that all of those GCabi lines had significantly higher stomatal conductance and larger stomatal apertures (Suppl. Fig. 1). All experiments were conducted using at least three randomly selected independent lines of homozygous T3 and T4 plants.

### 2.3. Confocal microscopy imaging

Images were acquired using the Olympus IX 81 inverted laser scanning confocal microscope (Fluoview 500; Olympus; Tokyo) equipped with a 488-nm argon ion laser and a 60 × 1.0 NA PlanApo water immersion objective. GFP was excited by 488-nm light and the emission was collected using a BA 505–525 filter. A BA 660 IF emission filter was used to observe chlorophyll autofluorescence. Confocal optical sections were obtained at 0.5-μm increments. The images were color-coded green for GFP and red for chlorophyll autofluorescence.

### 2.4. Stomatal measurements

Epidermal peels were soaked in ‘closure’ solutions, as described in Acharya et al. (2013), under a light intensity of ~150 μmol m−2 s−1 one hour after dawn. After 2 h, fusicoccin (Santa Cruz Biotechnology; Heidelberg, Germany) and ABA ((+)-cis, trans abscisic acid; Biosynth; Staad, Switzerland) were added to a final concentration of 10 µM. Solvents (ethanol and DMSO) were added to the control treatment at the same concentration. Fusicoccin is a fungal toxin that stimulates stomatal opening by activating the plasma membrane ATPase even in the presence of supplemental ABA. The fusicoccin treatment served as a positive control (Suppl. Fig. 2).

The stomatal apertures (Figs. 5 and 6), stomatal densities and stomatal indices of the plants were determined using the rapid imprinting technique described by Geisler and Sack (2002). This approach allowed us to reliably score hundreds of stomata from each treatment, each of which was sampled at the desired time. In brief, light-bodied vinylpolysiloxane dental resin (Heraeus-Kulzer; Hananu, Germany) was attached to the abaxial leaf side and then removed as soon as it had dried (1 min). The resin epidermal imprints were covered with nail polish, which was removed once it had dried. The nail-polish imprints were mirror images of the resin imprints. The nail-polish imprints were put onto microscope slides.

All stomata were photographed under a bright-field inverted microscope (1M7100; Zeiss; Jena, Germany) on which a Hitachi HV-D30 CCD camera (Hitachi; Tokyo, Japan) was mounted. Stomatal images were analyzed to determine aperture size using the ImageJ software (http://rsb.info.nih.gov/ij/). A microscopic ruler (Olympus; Tokyo, Japan) was used. Stomatal index was calculated as the number of stomata / (number of stomata + number of epidermal cells).

### 2.5. Measurements of whole-plant continuous canopy conductance

Whole-plant continuous canopy conductance (g_sc_) was measured using an array of load-cell lysimeters (Plantarray Gravimetric Prototype system, Plant-DiTech Ltd; Rehovot, Israel), as described by Halperin *et al.* (2016). GCabi and WT Arabidopsis plants were plated on kanamycin (50 mg mL–1) selection medium or antibiotic-free medium, respectively. After 3 weeks, the plantlets were transferred to 3.9-L pots (six plants per pot), which were kept in the greenhouse. Pots were covered with plastic wrap and gradually uncovered. *C. falcatum* ferns were planted directly into 3.9-L pots (one plant per pot).

The Arabidopsis plants were grown in a greenhouse under semi-controlled conditions of 26/12°C (day/night) and natural day length and light conditions in Rehovot, Israel during January and February of 2014. The ferns were grown in the greenhouse under a shade net and semi-controlled conditions of 27/18°C (day/night) and natural day length in Rehovot, Israel during May and June 2016. Daily measurements were conducted simultaneously for all of the plants in the array, so that all the plants were exposed to similar ambient conditions at each measurement point. Since no differences were observed between the three independent lines of GCabi used in previous experiments, only one line (GCabi9) was used in the lysimeter experiment, which allowed us to increase the number of replicates of GCabi. Each pot was placed on a temperature-compensated load cell. The soil surface surrounding each Arabidopsis plant was covered to prevent evaporation. The pots holding ferns were not sealed, due to the large area from which fronds were initiated. The output (weight) of the load cells was monitored every 10 s and 3-min average values were logged in a data-logger for further analysis. Whole-plant transpiration was calculated as a numerical derivative of the load-cell output following a data-smoothing process. The daily water loss rate was normalized to the total plant weight to determine the transpiration rate. Continuous whole-canopy conductance was calculated by dividing the whole-plant transpiration rate by the VPD.

### 2.6. Gas-exchange measurements

Leaves of plants that were 7 to 9 weeks old were excised just before dawn and immediately immersed (petiole-deep) in artificial xylem sap (AXS; 3 mM KNO_3_, 1 mM Ca(NO_3_)_2_, 1 mM MgSO_4_, 3 mM CaCl_2_, 0.25 mM NaH_2_PO_4_, 90 µM EDFC and a micromix of 0.0025 µM CuSO_4_*5H_2_O, 0.0025 µM H_2_MoO_4_, 0.01 µM MnSO_4_, 0.25 µM KCl, 0.125 µM H_3_BO_3_*3H_2_O, 0.01 µM ZnSO_4_*7 H_2_O). Cotton swabs were then used to smear leaves with 10 µM fusicoccin (Santa Cruz Biotechnology) dissolved in ethanol and diluted with AXS, or with AXS containing the same concentration of ethanol. The leaves were then kept in the growth chamber for 1 h to allow the smeared material to dry. Then, the leaves were put into a sealed transparent plastic box, in which they were exposed to elevated humidity, up to 94% (VPD = ~0. 15 kPa), for 2 h. From the beginning of the experiment, the boxes were kept in the lab under a light intensity of ~150 μmol m−2 s−1. The measurement data described below were collected using leaves from different boxes, in order to ensure a uniform, very humid starting point for all measurements

Gas-exchange measurements were taken using the LI-6400 portable gas-exchange system (LI-COR; Lincoln, NE, USA). In order to imprint leaves before and after the increase in VPD, pairs of leaves were prepared. At the beginning of each measurement, two leaves were taken from a box (VPD = ~0. 15 kPa), one leaf was immediately imprinted while the second leaf was placed in the LI-COR chamber for 20 min and then immediately imprinted. Measurements began 3 min after the leaf was placed in chamber, when the conditions in the chamber had. VPD was adjusted manually by adjusting the desiccant scrub flow during the 20 min (VPD = 0.93–1.07 kPa). The slope of the linear region of leaf response, from 9–17 min, was calculated. All measurements were taken between 10:00 and 15:00.

For the fern gas-exchange measurements, fronds were cut under water during the morning hours (8:00–8:30). From each frond, five leaflets (starting from the third leaflet from the top) were cut underwater and inserted into different Eppendorf tubes; the leaflets of each frond constituted a block. All treatments included AXS. Tubes that contained no DMSO were used to assess whether DMSO itself affected the gas exchange of *C. falcatum* (as was found in a previous experiment in which 0.4% DMSO was used). Blocks of the five 1.5-mL tubes were then put into hermetically sealed transparent plastic boxes and left under lights for 1 h. Following that hour, the boxes were opened for 5 min and measurement data was then collected.

### 2.7. Dark treatments

Two hours after dawn, well-watered whole plants were moved to darkness for 1 h. After that hour, g_s_ was measured using a leaf porometer (SC-1 Porometer; Decagon Devices, Inc., WA, USA). The plants were then moved back into the light (~150 μmol m−2 s−1) for an additional hour and g_s_ was then measured once again.

### 2.8. Stomatal conductance in drying soil

Stomatal conductance of 10- to 13-week-old plants was measured using a leaf porometer (SC-1 Porometer) and volumetric water content was measured with a Prochek probe (Decagon Devices). All measurements were carried out between 10:00 and 13:00.

### 2.9. Petiole-dip perfusion and leaf relative water content

Leaves excised before dawn were immediately immersed (petioles only, as shown in Fig. 3) in AXS and kept at close to zero VPD for 2 h (in a humid, transparent box), followed by 2 h of exposure to ambient VPD (~ 0.7 kPa) under light (125 µmol s-1 m-2). The duration and efficiency of the xylem-loading perfusion were confirmed in separate leaves under the same experimental conditions by following the red dye Safranin O (1% w/v) through the leaf veins (Sigma Cat. No. S2255, 1% w/w in AXS; Fig. 3A).

Relative water content (RWC) was measured as described by Sade *et al.* (2015). In short, leaf fresh weight (FW) was immediately recorded and leaves were then soaked for 8 h in 5 mM CaCl_2_ at room temperature in the dark, after which the turgid weight (TW) was recorded. Total dry weight (DW) was recorded after the leaves were dried at 70°C to a constant weight. RWC was calculated as (FW – DW / TW – DW) × 100.

### 2.10. Statistical analysis

Student’s *t*-test was used for comparisons of two means and the Tukey-Kramer test was used for comparisons of more than two means. Dunnett’s test was used for comparisons with the control. The Kruskall-Wallis non-parametric (one-way) test was used when the variance was not homogeneous. All analyses were done using JMP software (SAS; Cary, NC, USA).

## 3. Results

### 3.1. Responses of GCabi stomata to ABA, drought and darkness

After confirming the GC-specific expression of GFP under the KST promoter (GCGFP; Fig. 1A), we examined GCabi’s stomatal responses to ABA and drought. Initially, we examined the stomatal apertures of WT and GCabi epidermal peels that had been soaked in 10 µM ABA or 10 µM fusicoccin (a fungal toxin that stimulates stomatal opening). As expected, ABA caused a significant reduction in the stomatal apertures of the WT. However it had no significant effect on the stomatal aperture of GCabi (Fig. 2A), indicating that the quantity-dependent dominancy of *abi1-1* (Wu *et al.*, 2003) is maintained under the KST promotor. Moreover, under controlled conditions, the stomatal apertures of GCabi were significantly larger than those of the WT (5.22 + 0.06 and 4.66 + 0.08 µm, respectively). Fusicoccin led to the enlargement of stomatal apertures in both the WT and GCabi, resulting in similar wide-open apertures (>5.7 µm; Fig. 2A) among both sets of plants.

**Figure 1.**
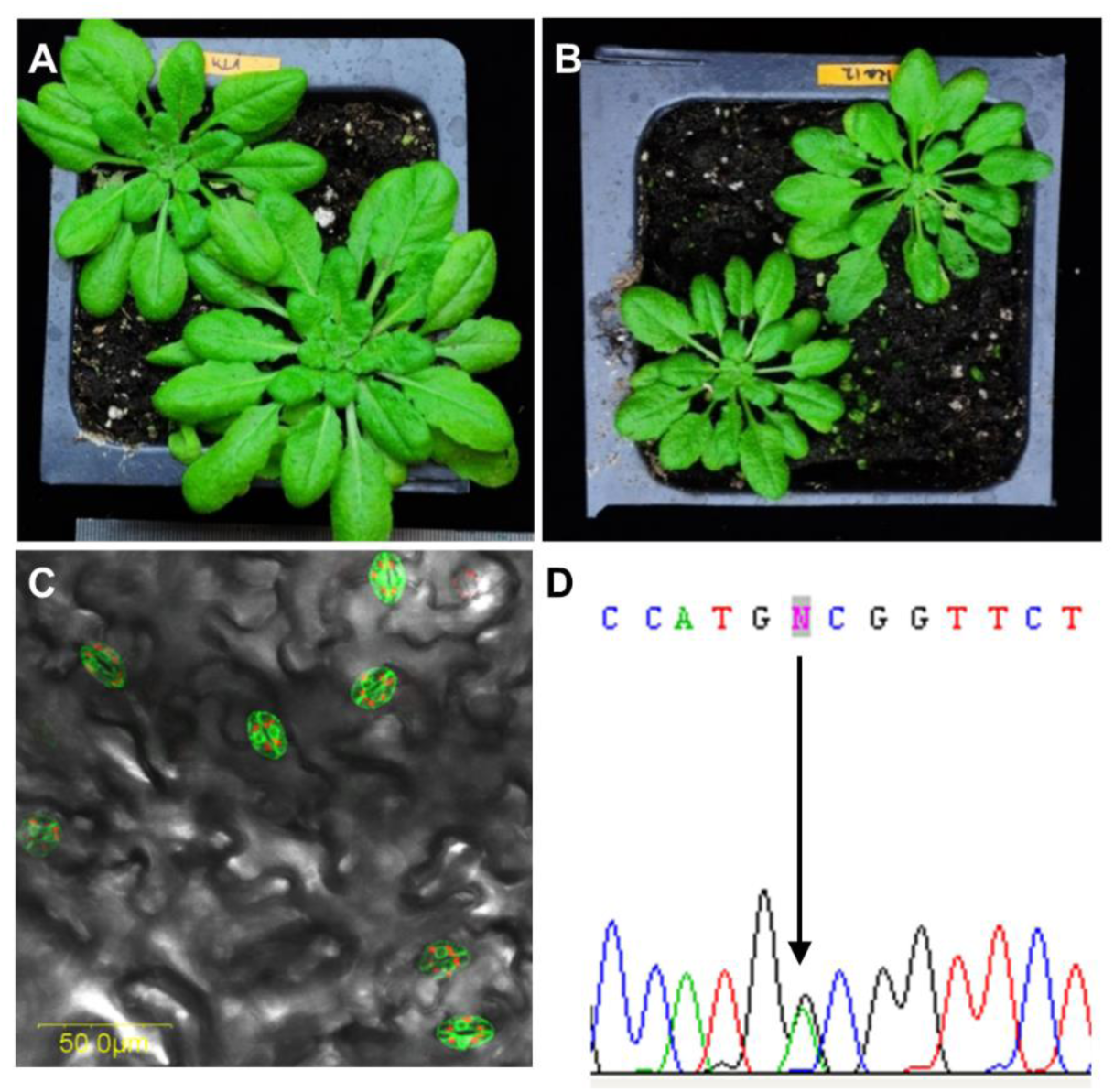
GCabi plants. Six-week-old WT (A) Colombia and (B) GCabi plants. (C) A fluorescent image (488-nm excitation; 520-nm emission) of a leaf expressing GFP under the KST promoter. (D) Sequence of KST:abi (GCabi) cDNA, the arrow points to the G→A mutation. This figure is available in colour at <u>*JXB online*</u>.

**Figure 2.**
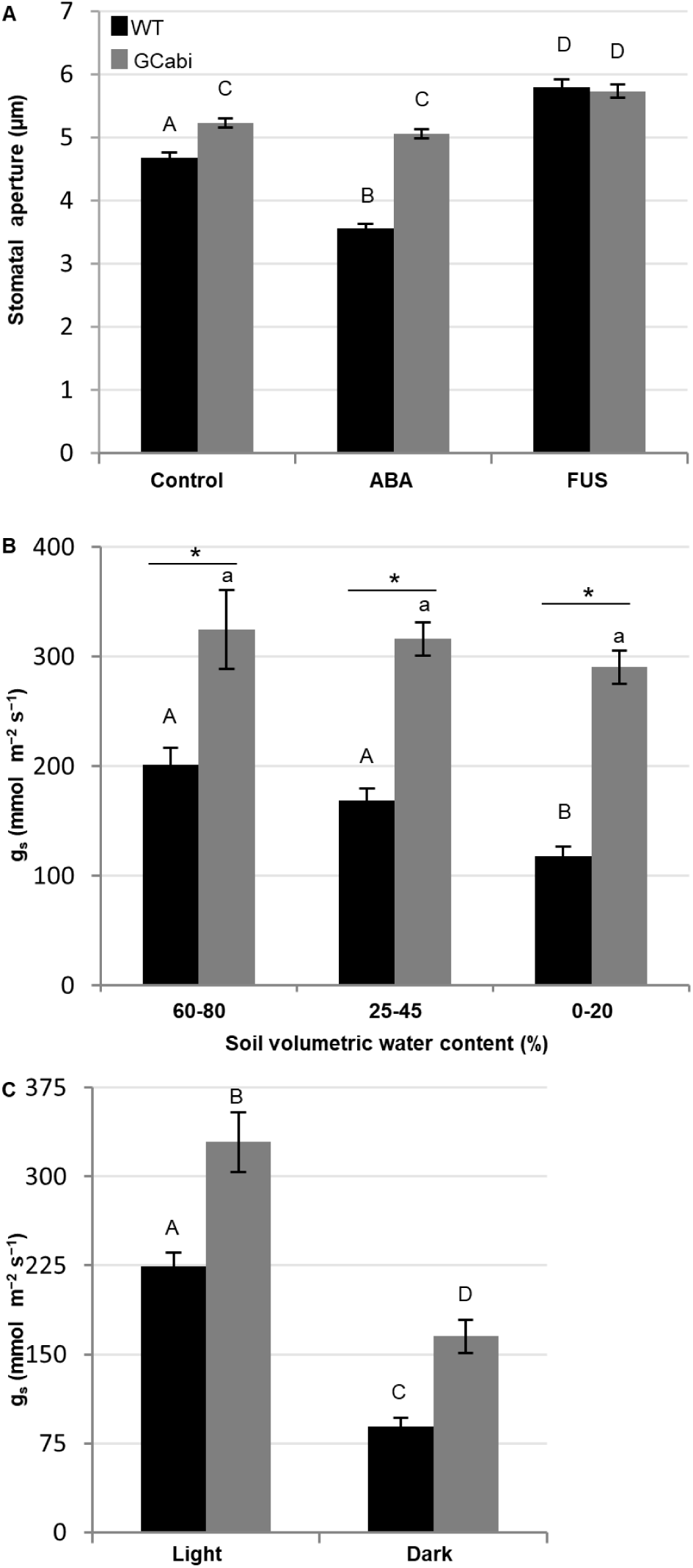
GCabi exhibits no significant stomatal response to external ABA or drought. (A) Stomatal apertures of WT (black bars) and GCabi (gray bars) epidermal peels directly exposed to 10 µM ABA or 10 µM fusicoccin. Data points are averages of at least three epidermal peels and represent a minimum of 160 stomata. (B) Stomatal conductance of 10-to 13-week-old plants in response to continuous drought and (C) stomatal conductance of 10-to 13-week-old plants exposed to light (125 µmol s-1 m-2) and then to 1 h darkness. Results are means + SE of at least 3 independent experiments and 3 independent lines of GCabi (for B, *n* = 25–100; for C, *n* = 63). (A, C) Different letters indicate a significant difference according to the Tukey-Kramer test (*P* < 0.05). (B) Different letters indicate a significant difference between treatments within the same line and asterisks indicate a significant difference between lines subjected to the same treatment, according to the Kruskall-Wallis non-parametric test (*P* < 0.05).

The GCabi plants exhibited no significant reduction in their stomatal conductance (g_s_) in response to reductions in soil volumetric water content and their stomatal conductance was significantly higher than that of the WT plants throughout the soil-drying treatment (Fig. 2B). The unimpaired responses of GCabi stomata to darkness and fusicoccin (Fig. 2C, A) indicate that these plants possess a functional stomatal-movement mechanism. The higher g_s_ and water-loss rate of the GCabi plants, as compared to the WT, led to the lower leaf relative water content (RWC) observed in detached leaves from those plants, in which minimal hydraulic resistance had been expected (Fig. 3). However, that was not the case for the whole GCabi plants, in which bulk flow from the roots was not disturbed. Two hours after leaves were excised and immersed (petiole-deep) in artificial xylem sap, the RWC of GCabi leaves was significantly lower than that of the untreated WT leaves (70.5% + 3.5 and 86.7% + 1.1, respectively) and not significantly different from that of the WT leaves that had been smeared with fusicoccin (77.5% + 1.1). This wilting may indicate that the high rate of water loss through stomata could not be compensated for by the leaf hydraulic conductance.

**Figure 3.**
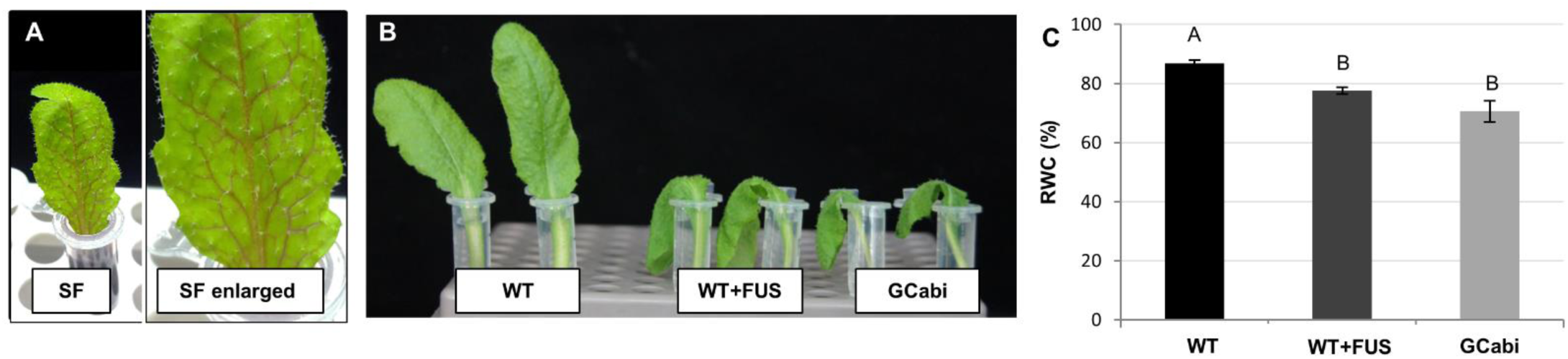
Perfusion of detached leaves via their petioles (petiole dip). The low-level stomatal regulation of GCabi leads to lower relative water content (RWC) in a manner similar to that observed among WT leaves smeared with fusicoccin (FUS). The efficacy of the petiole-dip perfusion (see Materials and methods) was confirmed by the fact that the xylem-borne dye spread throughout the leaf vasculature. WT leaves, WT leaves smeared with 10 µM fusicoccin and GCabi leaves were petiole-dipped in AXS without any safranin and (C) their relative water contents are shown. Results are means + SE of 3 independent experiments (*n* = 15). Three independent lines of GCabi plants were used. Different letters indicate a significant difference (Tukey-Kramer test, *P* < 0.05). This figure is available in color at *JXB online*.

### 3.2. GCabi plants exhibit a daily whole-plant canopy conductance pattern that is similar to that of the WT and different from that seen in the ferns

g_s_ is a dynamic parameter that changes over the course of the day in response to changes in environmental factors such as light and VPD. Since GCabi plants exhibit larger stomatal apertures and higher stomatal conductance, we were interested in monitoring their responses to daily changes in atmospheric conditions in the greenhouse. In order to measure the whole-plant canopy conductance (g_sc_) continuously among the GCabi plants and the control plants simultaneously, we used an array of lysimeters (see Materials and methods). Both WT and GCabi revealed similar patterns of g_sc_ in response to the natural changes in the environmental conditions in the greenhouse, which included an increase in g_sc_ during the early morning (when VPD is low and light levels are increasing), followed by a decline in g_sc_ as VPD increased, down to a steady-state during late morning and the middle of the day (Fig. 4A, B). Despite the similar g_sc_ patterns of GCabi and the WT, under well-irrigated conditions, GCabi exhibited significantly higher canopy conductance during most of the day (from 08:05 to 15:20; Fig. 4B).

**Figure 4.**
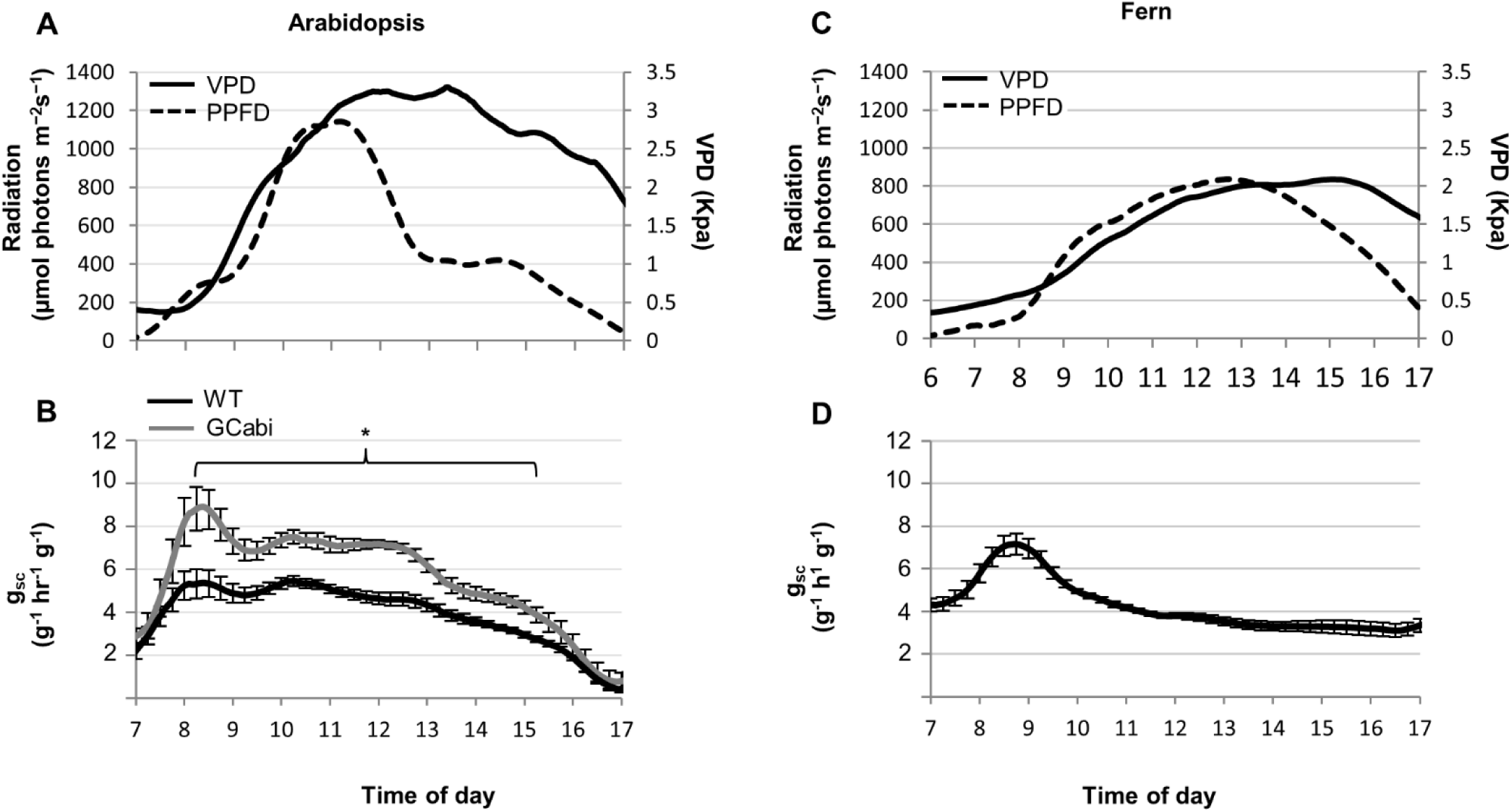
Daily pattern of whole-canopy stomatal conductance. GCabi and WT Arabidopsis plants and ferns were grown under well-irrigated greenhouse conditions. (A) Daily VPD (solid line) and light intensity (radiation, dashed line) for the Arabidopsis plants. (B) The whole-plant canopy stomatal conductance (gsc; g-1 h-1 unit plant weight-1; plant weight, g) of GCabi (gray) and WT (black) Arabidopsis plants.(C) Daily VPD (solid line) and light intensity (radiation, dashed line) for the ferns and (D) the whole-plant canopy stomatal conductance of the ferns. The relatively high basal level of fern g_sc_ is related to the fact that the soil surrounding those plants was not covered, due to the ferns’ dense growth habit (see Materials and methods). Curves show the means of 5 to 9 independent pots. Each pot included 6 plants; WT Arabidopsis plants (black, *n* = 45), GCabi plants (line GCabi9, gray, *n* = 25) and ferns (*n = 5*). The differences in ambient conditions (radiation and VPD) were due to the different growing seasons (winter for Arabidopsis and summer for fern). Data are shown as means ± SE. The asterisk indicates a significant difference between GCabi and WT according Student’s *t*-test (*P* < 0.005).

The ferns’ daily g_sc_ pattern was different from that observed for Arabidopsis, including an increase in g_sc_ during the early morning and a decrease down to the basal level as VPD increased during the early morning (Fig. 4C, D). Since fern insensitivity to ABA has been shown to be species- and growth condition-dependent (Hõrak *et al.*, 2017), we confirmed the insensitivity of *C. falcatum* to ABA in work with petiole-fed ABA (Suppl. Fig. 3). The similar responses of the g_sc_ of GCabi and the WT to changes in ambient conditions point to similar VPD-sensing in both types of plants or a stronger effect of some other signal such as light. Therefore, we decided to test the VPD-specific response of GC in a tightly controlled gas-exchange experiment.

### 3.3. GCabi plants exhibit a WT-like stomatal response to an increase in VPD

To study the role of ABA in the regulation of stomatal response to a sharp increase in VPD, we monitored changes in g_s_ over a period of 20 min, starting 3 min after leaves were transferred from low VPD conditions [0.15 kPa (high humidity)] to higher VPD conditions [1 kPa (lower humidity)], using the LI-COR 6400 chamber. Data were collected as soon as the chamber conditions stabilized (see Materials and methods). We also measured stomatal aperture before and after VPD was increased (Fig. 5). Fusicoccin treatment, which induces irreversible stomatal opening, was used as a positive control for non-sensitive open stomata. As expected, the sharp increase in VPD (from 0.15 to 1 kPa) resulted in a reduction in the g_s_ of the WT (Fig. 5A), which was correlated with a reduction in the stomatal apertures of those leaves (Fig. 5C). Interestingly, WT leaves that had been treated with fusicoccin also exhibited reduced stomatal conductance (Fig. 5A), yet with a significantly more moderate slope than that observed for the untreated WT (Fig. 5B) and a smaller, yet significant reduction in stomatal aperture (Fig. 5C). As before, the g_s_ and stomatal apertures of GCabi were significantly greater than those of the WT throughout the experiment (Fig. 5A, C). Nevertheless, the stomatal response patterns of GCabi to the jump in VPD, in terms of both g_s_ and stomatal aperture, were similar to the response patterns observed for the WT. The g_s_ graphs of the two sets of plants had the same slope (0.33 + 0.013 mmol m^−2^ s^−1^/min) and the two sets of plants also exhibited similar reductions in stomatal aperture (48% for the WT and 46% for GCabi; Fig. 5B).

**Figure 5.**
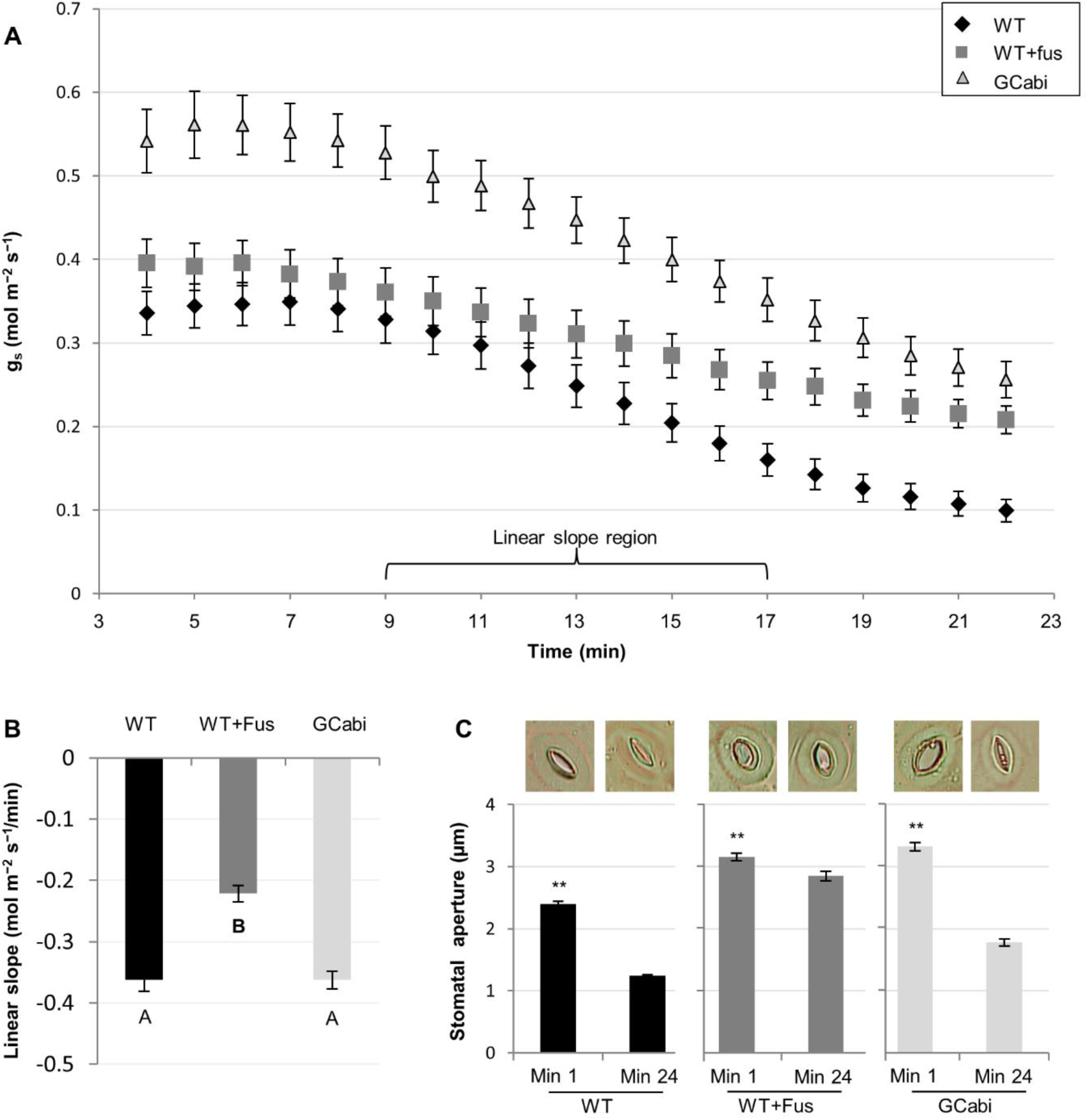
GCabi plants exhibit a WT-like stomatal response to a rapid increase in VPD. (A) Changes in stomatal conductance over time in response to an increase in VPD from 0.15 kPa to 1 kPa; WT (diamond), WT smeared with 10 µM fusicoccin (square) and GCabi (triangle). Measurements began 3 min after a leaf was placed in the gas-exchange chamber (see Material and methods). (B) The linear slope of g_s_ (9 to 17 min). (C) Bright-field microscopy images of stomatal imprints, measured on duplicate leaves exposed to a change in VPD at Minute 1 and Minute 24. The measurement data are also presented in bar graphs. The data shown in (A) and (B) are means of 20 leaves and at least 380 stomata for (C). Significant differences are indicated by letters (Tukey-Kramer test, *P* < 0.05) or by asterisks (*t*-test, *P* < 0.01).

### 3.4. Responses of GCabi and the WT to different VPD conditions

The higher g_s_ of the GCabi plants can be explained by their larger stomatal apertures (Figs. 2A, 5C) and their higher stomatal density. The fact that GCabi plants lose more water through transpiration raises the possibility that lack of leaf turgor may lead to their smaller-leaf phenotype (Fig. 6A) and, subsequently, to their higher stomatal density (Fig. 6B). Therefore, we grew the plants under ambient (1.4 kPa) and low VPD (0.2 kPa, with a transparent plastic lid kept over the growth tray to reduce transpiration and increase RWC). Indeed, the low VPD conditions restored the RWC of GCabi to the level observed for the WT leaves. That is, WT plants were able to preserve relatively high RWC under both high- and low-VPD conditions; whereas the RWC of GCabi decreased under ambient VPD conditions (Fig. 6C). Nevertheless, despite the RWC differences, GCabi leaf area did not change and remained smaller than that of the WT under both high and low VPD conditions (Fig. 6A). The small size of the GCabi leaves cannot fully explain GCabi’s higher stomatal density under higher VPD conditions. (WT stomata were 1.48 times larger; whereas GCabi stomata were 1.7 times denser). Therefore, stomatal index (number of stomata per epidermal cell) was measured as well. The GCabi stomatal index was higher than that of the WT under ambient conditions and decreased to match the unchanged stomatal index of the WT under low-VPD conditions (Fig. 6D). Interestingly, the total number of stomata per leaf for GCabi and WT was similar under both VPD growing conditions (Fig. 6E). In addition, the long-term, low-VPD conditions increased WT stomatal apertures to the level seen for GCabi. In contrast, GCabi stomatal aperture remained constant under the two VPD conditions (Fig. 6F).

**Figure 6.**
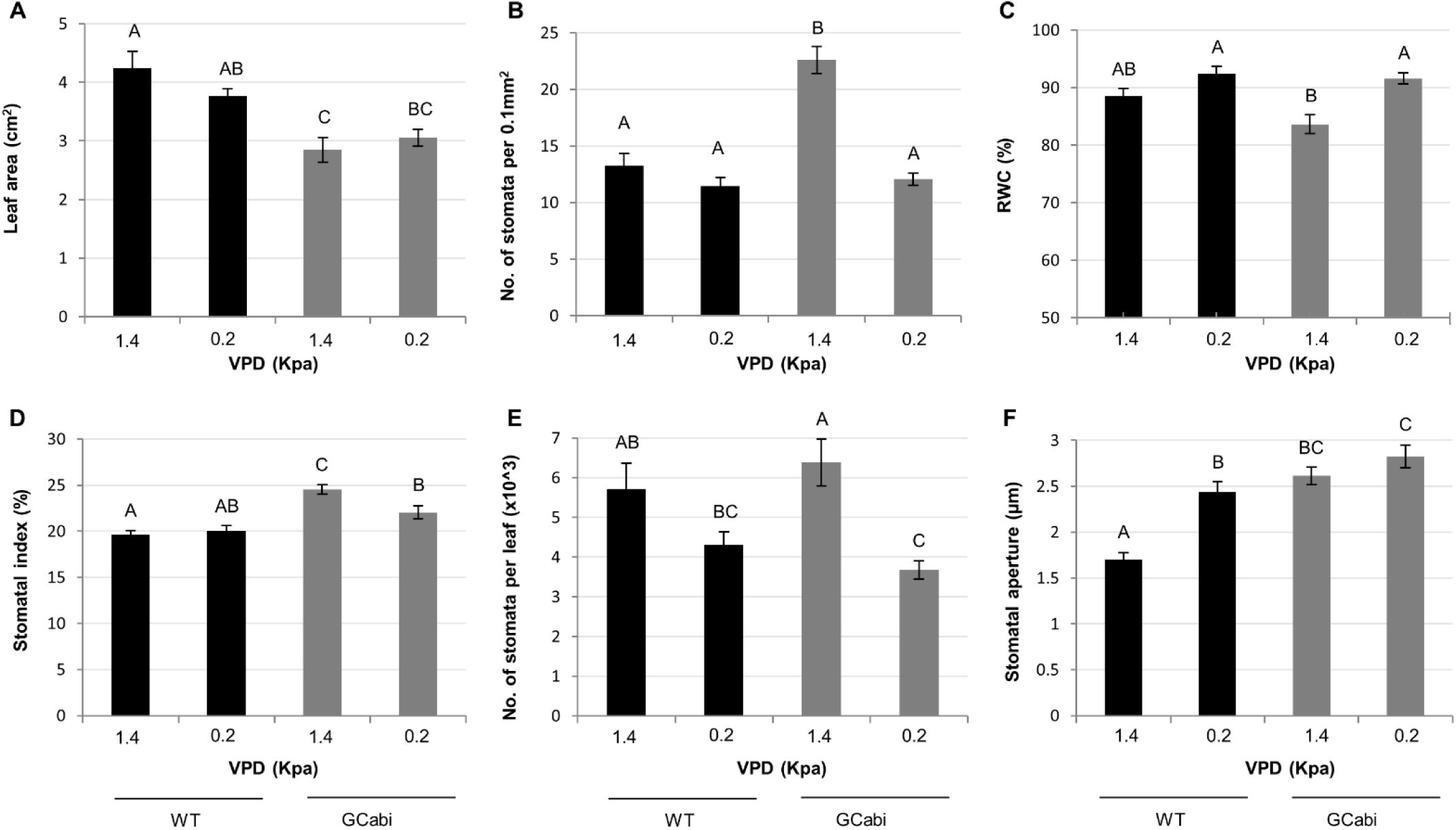
Stomatal characteristics of GCabi and WT Arabidopsis plants in response to ambient and low VPD conditions. Eight-week-old GCabi and WT plants were grown under ambient (1.4 kPa) or low (0.2 kPa) VPD conditions. (A) Leaf area; (B) stomatal density per 0.1 mm2 of leaf area; (C) leaf RWC; (D) stomatal index; (E) number of stomata per leaf and (F) stomatal aperture. Stomatal density, aperture and index were examined in 3 regions of 5 leaves from each treatment. For leaf area and RWC, *n* = 15. Results are means ± SE; different letters indicate a significant difference (Tukey-Kramer test, *P* < 0.05).

## 4. Discussion

### 4.1. ABA's role in regulating stomatal aperture under non-stressful conditions

The fact that the GC-specific ABA-insensitive plants (GCabi) had significantly larger stomatal apertures and greater stomatal and canopy conductance than the WT under well-irrigated conditions (Figs. 2, 4B, 5AC, 6F) supports the findings of previous studies, which have suggested that ABA plays a housekeeping role in limiting maximal stomatal apertures under non-stressful conditions (Kelly *et al.*, 2013; Pantin *et al.*, 2013*b*; Merilo *et al.*, 2017). Moreover, the fact that higher VPD caused a change in WT stomatal apertures, but not GCabi stomatal apertures (Fig. 6F) supports the theory that GC sensitivity to ABA plays a key role in that housekeeping role. The non-maximal aperture mediated by ABA has been hypothesized to play a role in one of the following optimization process: 1) improving plant water-use efficiency (WUE; Yoo *et al.*, 2009); 2) coordinating transpiration with photosynthesis (Kelly *et al.*, 2013); or 3) coordinating transpiration with vascular hydraulic limitations that may make the plant incapable of supporting the excessive transfer of water to transpiring leaves (Sack and Holbrook, 2006), leading to reduced leaf water potential (Shatil-Cohen *et al.*, 2011; Pantin *et al.*, 2013*a*). Lack of sufficient hydraulic conductivity can also explain the low RWC of the GCabi and WT + Fus detached leaves, as compared with the untreated WT (Fig. 3C), despite the fact that these leaves were submerged in solution and xylem-borne dye moved freely throughout each leaf (Fig. 3A, as well as Shatil-Cohen *et al.*, 2011, 2012).

Our results emphasize the fact that this stomatal aperture-limiting role of ABA is related specifically to the GC, as opposed to being a byproduct of the effect of ABA on hydraulic signals [i.e., vascular radial conductance or mesophyll water permeability, as demonstrated by Shatil-Cohen *et al.* (2011) and Pantin *et al.* (2013*a*)]. Such a hydraulic effect of a non-stomatal ABA response could have been involved in previously reported observations of whole-plant ABA-mutant lines and lines in which ABA production was limited to the GC (Bauer *et al.*, 2013; Merilo *et al.*, 2017) or phloem (Merilo *et al.*, 2017). To address that issue, we performed this work using a mutant in which the GC were the only cells insensitive to ABA. Indeed, Merilo *et al.* (2017) showed that the g_s_ and hysteresis of an ABA-deficient mutant were correlated with leaf ABA levels; whereas an ABA-insensitive mutant (whole-plant mutant; mutant *112458*) exhibited an altered response to change in VPD. That finding contrasts with the results of our work with a GC-specific ABA-insensitive mutant (which exhibited a pattern similar to that observed for the WT). As the main difference between the two ABA-insensitive plants is the sensing tissue (GC in our experiment and the whole plant in the *112458* mutant), it may be that some internal-tissue response to ABA was involved in GCabi’s response to VPD change, but not in that of the *112458* mutant, further emphasizing the importance of internal-tissue feedback for stomatal activity and the importance of GC ABA for determining maximal stomatal aperture.

### 4.2. Stomata–VPD relations and ABA's role in the passive-hydraulic g_s_ response

Typically, the daily pattern of g_s_ is strongly correlated with daily changes in VPD. This daily g_s_–VPD pattern is characterized by high g_s_ in the early morning when VPD is low and a decrease in g_s_ as VPD increases during morning hours, as the temperature rises and the relative humidity falls (Ullmann *et al.*, 1985; Raschke and Resemann, 1986; Brodribb and Holbrook, 2004; Kelly *et al.*, 2013; Halperin *et al.*, 2016; Fig. 3A, C). Obviously, this daily g_s_–VPD pattern is influenced by seasonal conditions, for example, higher maximal aperture and a slower rate of decrease are observed under the lower VPD levels typical of a rainy season (Brodribb and Holbrook, 2004). The impact of lower-VPD conditions on maximal stomatal aperture was also detected among our WT plants grown under 0.2 kPa VPD, which had stomatal apertures that were 65% larger than those of WT plants grown under a VPD of 1.4 kPa. In contrast, this phenomenon was not observed among the GCabi plants, whose stomatal apertures were wider and unaffected by the change in VPD (Fig. 6F). The fact that the daily g_sc_–VPD pattern of GCabi was similar to that of the WT, but with higher levels of g_sc_ throughout the day (Fig. 4A, B), supports the claim that ABA plays a housekeeping role in limiting the potential size of stomatal apertures under non-stressful conditions, as well as its nonfunctioning in the passive hydraulic g_s_ reduction (Assmann *et al.*, 2000; Merilo *et al.*, 2017). Measurement of the daily g_sc_–VPD pattern of the ABA-insensitive fern (*C. falcatum*; Fig. 4C, D, Suppl. Fig. 3) revealed a similarity with the Arabidopsis morning g_sc_ peak, which was followed by a decline (back to basal level) in g_sc_ by late morning. It was previously suggested that the stomatal closure-response of ferns to an increase in VPD could be a passive hydraulic response that does not involve ABA (Brodribb and McAdam, 2011; McAdam and Brodribb, 2014, 2015). Specifically, the ferns’ GC lose turgor as the VPD grows, resulting in stomatal closure that is not mediated by ABA. Conifers (which represent a phylogenetic midpoint between the fern and angiosperm clades) represent an intermediate stage in the development of ABA stomata regulation, in which ABA enhances stomatal closure under drought stress (Brodribb and McAdam, 2013), but is not involved in stomatal responses to VPD (McAdam and Brodribb, 2015). The Arabidopsis ABA-independent response to an increase in VPD was replicated in a tightly controlled gas-exchange chamber, in which the patterns of stomatal aperture and g_s_ responses of GCabi leaves were similar to those observed for the WT (Fig. 4).

This ABA-independent response of GC to VPD may be due to the passive-hydraulic response mechanism, which may be attributed to either ancestral regulation that has remained significant in some angiosperm species, including Arabidopsis (McAdam and Brodribb, 2015), or a mechanism that regulates the GC response to a new steady state in bulk leaf turgor (i.e., the new balance between pressures of the GC and epidermal cells; Glinka and Aviv, 1971; Zait *et al.*, 2017). The second possibility could explain GCabi’s larger apertures under ambient conditions (operating close to turgor loss point; Fig. 2), so that GC turgor dominates epidermal pressure, as in a "continuous wrong-way response." Yet, GCabi stomatal aperture was unchanged when RWC increased (i.e., higher turgor, low VPD), weakening that argument (Fig. 5B, F). In addition, differences in the balance of pressures between the epidermis and GC are expected to be reflected in stomatal-closure dynamics that differ from those observed for the WT control, such as those seen for the fusicoccin-treated WT leaves, but not for GCabi (Fig. 5).

Alternatively, the ABA-independent response of GC to VPD could be due to a physical, as yet unknown parameter that co-varies with transpiration, so that the GC sense changes in the flux of water through the stomate (Mott, 1991; Assmann *et al.*, 2000). In that case, both the perception of changes in the transpiration rate and the mechanism by which that signal would be transduced remain unclear. OST1, a protein kinase active downstream of ABA, might be involved in such an ABA-independent response (Yoshida *et al.*, 2006; Merilo *et al.*, 2017).

### 4.3. ABA-independent and ABA-dependent responses to VPD

In light of the evidence presented above, it seems that (at least in Arabidopsis) the stomatal response to a sharp increase in VPD involves three elements: leaf hydraulic status, an ABA-independent mechanism and an ABA-dependent mechanism. A possible explanation that includes all three of these elements could be that ABA is not the initial cause of stomata closure, but rather a consequence of that closure. According to the above rationale, we suggest that while ABA became more and more dominant in stomatal regulation over the course of evolution (providing vascular plants with greater plasticity and helping them to adapt to new environments; Brodribb and McAdam, 2011; McAdam and Brodribb, 2015; Negin and Moshelion, 2017), angiosperms did not entirely lose their passive hydraulic stomatal-response mechanism. Nevertheless, the (symplastic) isolation of the GC from epidermal cells in the leaves of angiosperms (Kong *et al.*, 2012; Sager and Lee, 2014) limits that hydraulic response, as compared to the hydraulic response observed in ferns. This “hydraulic independency” allows for larger stomatal apertures, but also means that active turgor loss is required to reduce stomatal aperture beyond the initial hydraulic passive response. We hypothesized that the potential advantage of this combined strategy is the flexibility to have two modes of action: 1) high WUE with high stomatal conductance (i.e., enabling high CO_2_ intake) during low VPD periods and 2) the ability to keep stomata slightly open during periods of higher VPD, even at the price of lower WUE (risk-taking anisohydric behavior; Negin and Moshelion, 2017; Tardieu and Simonneau, 1998). In this hypothetical biphasic model, a passive hydraulic response triggers an active ABA-dependent response and ABA enables the optimal maximal g_s_ for each phase, corresponding to ambient conditions (i.e., soil water content and VPD).

The main role of ABA in the evolution of the angiosperms may have been in the adjustment of stomatal opening, as opposed to the common understanding of ABA as the stomatal-closing phytohormone. Accordingly, we can also explain angiosperms’ relative long phase of steady-state g_s_ (GC of both WT and GCabi maintained some turgor) during late morning and midday (Ullmann *et al.*, 1985; Raschke and Resemann, 1986; Brodribb and Holbrook, 2004; Kelly *et al.*, 2013; Halperin *et al.*, 2016; Fig. 4B) and the lag in the rate at which stomata open following a sharp increase in VPD, as compared to the rate at which they close (McAdam and Brodribb, 2015, 2016; Merilo *et al.*, 2017).

### 4.4. ABA’s role in adaptation to ambient VPD

The goal of growing plants under low VPD was to reduce their water loss and increase their RWC. In addition, this experiment uncovered some interesting long-term effects of VPD on stomatal development and the stomatal response to ABA. Low-VPD conditions increased the size of WT apertures, but did not affect the large stomatal apertures of the GCabi plants (Fig. 6F). This is in agreement with previous studies that have shown that VPD conditions (1 to 4 days of exposure) affect stomatal aperture (Fanourakis *et al.*, 2011; Aliniaeifard *et al.*, 2014; Carvalho *et al.*, 2015), the quantity of leaf ABA (Rezaei Nejad and van Meeteren, 2008; Arve *et al.*, 2013; Giday *et al.*, 2014) and stomatal sensitivity to ABA (Rezaei Nejad and van Meeteren, 2008; Aliniaeifard and Van Meeteren, 2013; Pantin *et al.*, 2013*b*; Arve *et al.*, 2014; Giday *et al.*, 2014), which is reversed by the application of ABA (Fanourakis *et al.*, 2011; Aliniaeifard *et al.*, 2014) or air movement (Carvalho *et al.*, 2015) during the low-VPD period. Growing GCabi plants under constant low-VPD conditions did not increase their leaf area, despite the observed increase in their RWC (Fig. 6A, C). This suggests that lack of turgor (Fig. 3B, 6C) is not the main cause of the small size of GCabi leaves and that the higher stomatal density of GCabi (Fig. 6B) is at least partially due to developmental modification, as seen in their higher stomatal index (Fig. 6D). Expressing *abi1-1* under a GC-specific promoter was sufficient to increase stomatal density relative to that observed for *abi1-1* mutants (Tanaka *et al.*, 2013). This explains why the g_s_ of the GCabi leaves was higher than that of WT leaves that were treated with fusicoccin, despite their similar stomatal apertures (Fig. 5A, C). Moreover, the fact that the promoter used to construct the GCabi plants is GC-specific (Müller-Röber *et al.*, 1995; Kelly *et al.*, 2013; Sade *et al.*, 2014; Fig. 1) and likely activated late in or even after differentiation suggests that ABA may have an indirect effect on stomatal proliferation through stomatal aperture and the transpiration rate (Lake and Woodward, 2008), in addition to the direct effect suggested by Tanaka *et al.* (2013). The greater stomatal density of GCabi was VPD-dependent and was reduced when those plants were grown under low-VPD conditions (Fig. 6B). This contrasts with the findings of previous studies, which suggested that low RH (high VPD; Tricker *et al.*, 2012; Chater *et al.*, 2014; Carvalho *et al.*, 2015) and/or ABA (Tanaka *et al.*, 2013) suppress stomatal proliferation. In our experiment, reducing the VPD increased the size of stomatal apertures (Fig. 6F) and, subsequently, WT g_s_ (Fanourakis *et al.*, 2011; Arve *et al.*, 2013; Aliniaeifard *et al.*, 2014). In contrast, the stomatal apertures of GCabi, which were large to begin with, had lower g_s_ under lower VPD conditions, which could annul the transduction of stress signals, reducing GCabi’s stomatal density and stomatal index to WT levels. The long-term outcome of these conditions resulted in the developmental changes observed, possibly balancing the loss of water through stomata with stomatal proliferation (Chater *et al.*, 2014).

### 4.5. Conclusion

In this study, we describe a biphasic GC response to ABA. We demonstrate the importance of GC ABA for restricting stomatal apertures under well-irrigated conditions, in contrast to its insignificance in the immediate GC response to changes in VPD. In addition, we demonstrate that GC-specific ABA plays an indirect role in stomatal proliferation.

We summarize this study with a daily VPD–g_s_ response-curve hypothesis (Fig. 7). This dynamic response-curve hypothesis is based on the notion that stomatal aperture always reflects the sum of signals sensed by the GC (e.g., light, CO_2_ and ABA). The VPD signal has a special dual effect as it is the physical force that drives transpiration and also serves as a (direct or indirect) closing signal. Hence, under the natural dynamic pattern of daily signals, a typical g_s_ curve of a well-irrigated plant is expected to show the following pattern (as illustrated in Fig. 7): The first daylight initiates stomatal opening. At that point, a continum between the leaf substomatal cuvity and the atmosphere (i.e., VPD) is established (i.e., the opening of stomata initiates the soil-plant-atmosphere continuum, which was blocked while the stomata were closed). At this early hour, VPD is at its lowest level and stomatal apertures are at their largest. As the temperature rises, VPD increases gradually, which leads to proportional water flux through the stomata that causes a passive hydraulic reduction in g_s_. In addition, the passive hydraulic response triggers corresponding ABA synthesis within minutes (McAdam and Brodribb, 2016; Sussmilch *et al*., 2017). The amount of ABA produced and GC sensitivity to ABA restrict the size of stomatal apertures, to keep g_s_, at a steady-state level that is appropriate for the prevailing ambient conditions.

**Figure 7.**
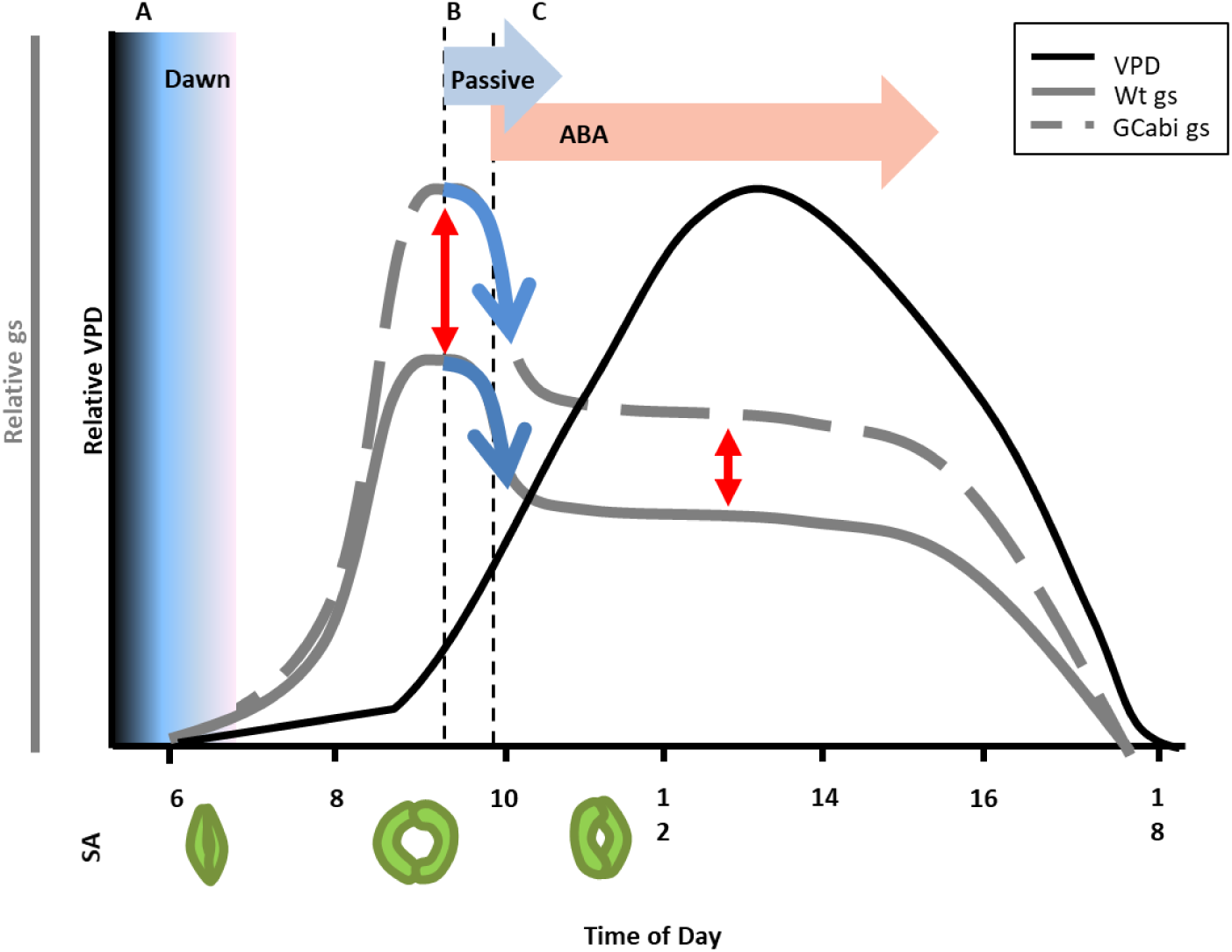
Our hypothetical biphasic stomatal VPD-sensing model. This model suggests that under well-irrigated conditions (A) the stomatal conductance (g_s_) of the WT (solid gray line) and GCabi (dashed gray line) generally increases rapidly, beginning at the first light at dawn, when VPD is low (black line), and maximal g_s_ (and maximal stomatal aperture, SA, vertical red arrow) is reached during the morning, in coordination with the sum of signals perceived by the GC, including the basal ABA level. (B) The increasing VPD induces a higher rate of transpiration, which triggers a reduction in SA and g_s_ via a passive-hydraulic, ABA-independent mechanism (blue arrows). (C) The passive-hydraulic signal induces the synthesis of ABA (horizontal red arrow), triggering the start of an ABA-dependent phase, which regulates the steady-state g_s_ and SA throughout the middle of the day and the afternoon (vertical red arrow). This VPD–ABA synthesis feedback may serve as a regulatory mechanism that enables the plant to optimize its SA under the prevailing VPD conditions.

## Abbreviations

(ABA): Abscisic acid

(AXS): artificial xylem sap

(GC): guard cells

(RWC): relative water content

(g_s_): stomatal conductance

(VPD): vapor pressure deficit

(WUE): water-use efficiency

## Supplementary Figures

**Fig. S1.** Stomatal aperture and g_s_ of all GCabi lines is compare to WT.

**Fig. S2.** Fusicoccin inhibits the effect of ABA on stomatal aperture in the WT.

**Fig. S3.** *Cyrtomium falcatum’*s insensitivity to ABA.

**Fig. S4**. Photo of representative WT and GCabi Arabidopsis plants grown under ambient and low VPD conditions.

## Acknowledgments

We thank Prof. Sarah Assmann for her knowledgeable remarks regarding the use of the *abi1-1* mutant gene and Prof. Dizza Bursztyn for her assistance with the statistical analysis.

